# Identification and characterization of retro-DNAs, a new type of retrotransposons originated from DNA transposons, in primate genomes

**DOI:** 10.1101/2020.03.19.999144

**Authors:** Wanxiangfu Tang, Ping Liang

**Affiliations:** Department of Biological Sciences, Brock University, 1812 Sir Isaac Brock Way, St. Catharines, Ontario, Canada L2S 3A1

**Keywords:** Primates, DNA transposons, Retrotransposons, Retro-DNA, Target-primed reverse transcription

## Abstract

Mobile elements (MEs) can be divided into two major classes based on their transposition mechanisms as retrotransposons and DNA transposons. DNA transposons move in the genomes directly in the form of DNA in a cut-and-paste style, while retrotransposons utilize an RNA-intermediate to transpose in a “copy-and-paste” fashion. In addition to the <u>t</u>arget <u>s</u>ite <u>d</u>uplications (TSDs), a hallmark of transposition shared by both classes, the DNA transposons also carry <u>t</u>erminal <u>i</u>nverted <u>r</u>epeats (TIRs). DNA transposons constitute ~3% of primate genomes and they are thought to be inactive in the recent primate genomes since ~37My ago despite their success during early primate evolution. Retrotransposons can be further divided into Long Terminal Repeat retrotransposons (LTRs), which are characterized by the presence of LTRs at the two ends, and non-LTRs, which lack LTRs. In the primate genomes, LTRs constitute ~9% of genomes and have a low level of ongoing activity, while non-LTR retrotransposons represent the major types of MEs, contributing to ~37% of the genomes with some members being very young and currently active in retrotransposition. The four known types of non-LTR retrotransposons include LINEs, SINEs, SVAs, and processed pseudogenes, all characterized by the presence of a polyA tail and TSDs, which mostly range from 8 to 15 bp in length. All non-LTR retrotransposons are known to utilize the L1-based target-primed reverse transcription (TPRT) machineries for retrotransposition. In this study, we report a new type of non-LTR retrotransposon, which we named as retro-DNAs, to represent DNA transposons by sequence but non-LTR retrotransposons by the transposition mechanism in the recent primate genomes. By using a bioinformatics comparative genomics approach, we identified a total of 1,750 retro-DNAs, which represent 748 unique insertion events in the human genome and nine non-human primate genomes from the ape and monkey groups. These retro-DNAs, mostly as fragments of full-length DNA transposons, carry no TIRs but longer TSDs with ~23.5% also carrying a polyA tail and with their insertion site motifs and TSD length pattern characteristic of non-LTR retrotransposons. These features suggest that these retro-DNAs are DNA transposon sequences likely mobilized by the TPRT mechanism. Further, at least 40% of these retro-DNAs locate to genic regions, presenting significant potentials for impacting gene function. More interestingly, some retro-DNAs, as well as their parent sites, show certain levels of current transcriptional expression, suggesting that they have the potential to create more retro-DNAs in the current primate genomes. The identification of retro-DNAs, despite small in number, reveals a new mechanism in propagating the DNA transposons sequences in the primate genomes with the absence of canonical DNA transposon activity. It also suggests that the L1 TPRT machinery may have the ability to retrotranspose a wider variety of DNA sequences than what we currently know.

## Introduction

Mobile elements (MEs), also known as transposable elements, as a whole constitute significant proportions of the genomes for most higher organisms, being around 50% in primate genomes (Carbone et al. 2014; Chimpanzee Sequencing and Analysis 2005; Cordaux and Batzer 2009b; Deininger et al. 2003; Higashino et al. 2012; Lander et al. 2001; Locke et al. 2011; Rhesus Macaque Genome Sequencing and Analysis et al. 2007; Scally et al. 2012; Tang and Liang 2019; Warren et al. 2015; Yan et al. 2011). MEs are defined as genomic sequences capable of changing locations or making copies into other locations within genomes. Despite being initially considered junk DNA, research from the last few decades has demonstrated that MEs have made significant contributions to genome evolution and they can impact gene function via a variety of mechanisms. These mechanisms include, but are not limited to, generation of insertional mutations and genomic instability, creation of new genes and splicing isoforms, exon shuffling, and alteration of gene expression and epigenetic regulation (Callinan et al. 2005; Chuong et al. 2016; Feschotte and Pritham 2007; J. S. Han et al. 2004; K. Han et al. 2005; K. Han et al. 2007; Konkel and Batzer 2010; Mita and Boeke 2016; Quinn and Bubb 2014; Sen et al. 2006; Symer et al. 2002; Szak et al. 2003; Wheelan et al. 2005). Additionally, MEs by germline or somatic insertions can contribute to genetic diseases in humans (see reviews by Anwar et al. 2017; Goodier 2016).

Based on the type of their transposition intermediates, MEs can be divided into two major classes: the Class I MEs or retrotransposons, which utilize an RNA-intermediate to transpose in a “copy-and-paste” fashion, and the Class II MEs or DNA transposons, which utilize a DNA-intermediate to transpose in a “cut-and-paste” style. Despite both having <u>t</u>arget <u>s</u>ite <u>d</u>uplications (TSDs), the two ME classes differ in their sequence characteristics, not only in their actual sequences, but also in TSD length and whether there are <u>t</u>erminal <u>i</u>nverted <u>r</u>epeats (TIRs) and polyA tail sequence, etc. (Feschotte and Pritham 2007; Pace Ii and Feschotte 2007; Smit and Riggs 1996).

Retrotransposons represent the majority of MEs in primate genomes, owing to their “copy-and-paste” style transposition, which results in direct copy number increase. In this process, a retrotransposon is first transcribed into RNA and then reverse transcribed into DNA as a new copy inserting into a new location in the genome (Kazazian and Goodier 2002). Retrotransposons can be divided into two major subtypes: the LTR and non-LTR retrotransposons, with the former carrying long terminal repeats (LTRs) that are absent from the latter, while the latter mostly have a polyA tail (Cordaux and Batzer 2009b; Deininger et al. 2003). In primates, LTR retrotransposons, mainly as endogenous retrovirus (ERVs) originated from retrovirus affecting and integrating into the germline genomes at various times during primate evolution, constitute ~9.0% of the genomes (Buzdin et al. 2003; Hughes and Coffin 2004; J. M. Kim et al. 1998; Lapuk et al. 1999). In comparison, non-LTR retrotransposons, as the most successful MEs in primate genomes, contribute to more than 35% of the genomes and more than 80% of all MEs (Tang and Liang 2019). By sequence features, the currently known non-LTR MEs belong to four subclasses, including Short-INterspersed Elements (SINEs), Long-INterspersed Elements (LINEs), SINE-R/VNTR/Alu (SVA), and processed pseudogenes (i.e. retro-copies of mRNAs) (Cordaux and Batzer 2009a; Ding et al. 2006; Kazazian and Moran 1998; Kazazian 2000; Ostertag and Kazazian 2001; Raiz et al. 2012). All subclasses of non-LTR retrotransposons, despite having many differences, such as size, sequencing feature, and coding capacity, share the common property of having a 3’ polyA tail and the use of <u>t</u>arget-<u>p</u>rime <u>r</u>everse <u>t</u>ranscription (TPRT) mechanism for retrotransposition (Goodier 2016; Ostertag and Kazazian 2001).

LINE-1s (L1s), being the only subfamily of autonomous non-LTR retrotransposons in the primate genomes, provide the TPRT machinery for all other non-LTR retrotransposons, which are considered non-autonomous for transposition (Cost and Boeke 1998; Goodier 2016; Jurka 1997; Mita and Boeke 2016; Tang et al. 2018; Xing et al. 2006). A functional L1, which is ~6,000 bp long, consists of an internal RNA polymerase II promoter, two open reading frames (ORF1 and ORF2) and a polyadenylation signal followed by a polyA tail (Kazazian and Goodier 2002). The ORF1 gene encodes an RNA-binding protein and ORF2 encodes a protein with endonuclease and reverse transcriptase activity (Cost et al. 2002; Feng et al. 1996; Goodier 2016; Kazazian and Goodier 2002; Martin 2006). Several studies have shown that Alus, L1s, and SVAs have an identical core sequence motif of “TT/AAAA” for their insertion sites, confirming that all non-LTR retrotransposition use the same TPRP mechanism (Cost and Boeke 1998; Jurka 1997; Tang et al. 2018; Wang et al. 2006).

In contrast to retrotransposons, DNA transposons, initially known as the “jumping genes,” move in genomes using a transposase encoded by autonomous copies (Deininger et al. 2003). Ten out of the twelve DNA transposon superfamilies are known to excise themselves out from their original locations as double-stranded DNA and move to new sites in the genome, which leads to no direct change in copy numbers (Feschotte and Pritham 2007; Pace Ii and Feschotte 2007). Two of the superfamilies, *Helitrons* and *Mavericks*, transpose through non-canonical mechanisms by utilizing a single-stranded DNA as intermediate, which leads to a “copy-and-paste” style (Feschotte and Pritham 2007; Kapitonov and Jurka 2001; Pritham et al. 2007). The ten “cut-and-paste” DNA transposon superfamilies, as well as *Mavericks*, have the presence of TIRs and TSDs, while *Helitrons* is the only superfamily with neither TIRs nor TSDs, owing to its rolling-circle mechanism (Feschotte and Pritham 2007; Kapitonov and Jurka 2001). In addition to these aforementioned DNA transposons, there is another group of DNA transposons named <u>m</u>iniature <u>i</u>nverted-repeat <u>t</u>ransposable <u>e</u>lement (MITEs) characterized by the presence of both TSDs and TIRs yet lacking the coding capacity for the transposase (Zhang et al. 2000). By using DNA transpose encoded by other autonomous DNA transposons, these non-autonomous, short (50-600bp) MITE entries can transpose in the host genome (Feschotte et al. 2003; Feschotte and Pritham 2007).

Past studies on MEs in the primate genomes have been mainly focused on the retrotransposons due to their significant contribution to the genome and their active contribution to inter-and intra-species genetic variations as lineage-specific or species-specific MEs driven by young and active members (Ahmed et al. 2013; Battilana et al. 2006; Ewing and Kazazian 2011; Liang and Tang 2012; Stewart et al. 2011). Selected full L1 sequences from the human genome have been shown to have retrotransposition activity in vitro and in vivo (Coufal et al. 2009; Gilbert et al. 2005; Kopera et al. 2016; Moran 1999). In contrary, DNA transposons have been considered inactive in the current primate genomes and have received very little research attention. Lander et al. in their initial human genome analysis concluded that there was no evidence for DNA transposon activity during the past 50 My (Lander et al. 2001), while a later study suggested that DNA transposons had been highly active during the early part of primate evolution till ~37Mya (Pace Ii and Feschotte 2007). There have been very few, if any, published reports for lineage-specific or species-specific DNA transposons in primate genomes. However, in our recent comparative analysis of species-specific MEs in eight primates from the *Hominidae* and the *Cercopithecidae* families, in addition to the identification of 228,450 species-specific retrotransposons (Tang and Liang 2019), we also identified a total of 2,405 DNA transposons which are also species-specific that were not included in our report. As part of efforts to understand the mechanisms underlying these species-specific DNA transposons, we report in this study a new type of non-LTR retrotransposons derived from DNA transposons. These DNA transposons share sequence features characteristic of L1-based retrotransposons, and we therefore name them as retro-DNAs, adding them as the fifth subclass of non-LTR retrotransposons after LINEs, SINEs, SVAs, and processed pseudogenes.

## Materials and Methods

### Sources of primate genome sequences

In this study, we chose to use ten primate genomes including human, among which eight genomes were inluded in our previous study for identifying species-specific MEs in primates (Tang and Liang 2019). These 10 primate species include human (GRCh38/UCSC hg38), chimpanzee (May 2016, CSAC Pan_troglodytes-3.0/panTro5), gorilla (Dec 2014, NCBI project 31265/gorGor4.1), orangutan (Jul. 2007, WUSTL version Pongo_albelii-2.0.2/ponAbe2), gibbon (Oct. 2012 GGSC Nleu3.0/nomLeu3.0), green monkey (Mar. 2014 VGC Chlorocebus_sabeus-1.1/chlSab2), crab-eating macaque (Jun. 2013 WashU Macaca_fascicularis_5.0/macFas5), rhesus monkey (Nov. 2015 BCM Mmul_8.0.1/rheMac8), baboon (Anubis) (Mar. 2012 Baylor Panu_2.0/papAnu2), and marmoset (Mar. 2009 WUGSC 3.2/calJac3). All genome sequences in fasta format and the RepeatMasker annotation files were downloaded from the UCSC genomic website (http://genome.ucsc.edu) onto our local servers for in-house analyses. We have used the most recent genome versions available on the UCSC genome browser site in all cases except for gorilla. For the gorilla genome, there is a newer version (Mar. 2016, GSMRT3/gorGor5) available, but it was not scaffolded into chromosomes, making it difficult to be used for our purpose.

### LiftOver overchain file generation

A total of 90 liftOver chain files were needed for all possible pair-wise comparisons of the ten genomes used in this study. These files contain the information linking the orthologous positions in a pair of genomes based on lastZ alignment (Harris 2007). Twenty-two of these were available and downloaded from the UCSC genome browser site, and another 34 liftOver chain files were generated using a modified version of UCSC pipeline RunLastzChain (http://genome.ucsc.edu) from a previous study (Tang and Liang 2019). The remaining 36 liftOver chain files were newly generated using the same pipeline.

### Identification of DNA transposons with diallelic status in the ten primate genomes

#### Pre-processing of DNA transposon

The starting list of DNA transposons in each primate genome was obtained based on the RepeatMasker ME annotation data from the UCSC website (https://genome.ucsc.edu). As previously described, we performed a pre-processing to integrate the ME fragments annotated by RepeatMasker back to ME sequences representing the original transposition events (Tang et al. 2018).

#### Identification of DNA transposons with diallelic status

We modified a previously reported bioinformatics comparative genomics approach (Tang et al. 2018) to identify diallelic DNA transposons (da-DNAs) that have the presence of both the insertion and pre-integration alleles in the ten primate genomes. Briefly, this pipeline uses a robust multi-way computational comparative genomic approach to determine the presence/absence status of a ME among a group of genomes by using both the whole chromosome alignment-based liftOver tool and the local sequence alignment-based BLAT tool (Hinrichs et al. 2006; Kent 2002). The sequences of a DNA transposon at the insertion site and its two flanking regions in a genome were compared to the sequences of the orthologous regions available in all other genomes. If a DNA transposon is absent from the orthologous regions of any of the other nine genomes not due to the existence of a sequence gap, it is selected as a potential candidate for a da-DNA subject to further analyses.

### Identification of retro-DNAs

#### Identification of TSDs and TIRs

For the candidate entries from the previous step, using in-house PERL scripts as described previously (Tang et al. 2018), we performed identification of the TSDs. Additionally, we have modified our scripts to identify the TIRs, which are the hallmarks of all cut-and-paste transposons except for *Helitrons* (Feschotte and Pritham 2007). da-DNA entries without identifiable TSDs or TSD length < 8 bp, as well as entries with identifiable TIRs, were excluded from further analysis. The 8 bp TSD length cutoff was chosen based on our observation for human-specific retrotransposons that 95% of identified TSDs are at least 8 bp long (Tang et al. 2018). Additionally, we used *MiteFinderII*, a tool designed to identify MIMEs (Hu et al. 2018), to verify that none of our candidate entries contain TIRs.

#### Filtering against retrotransposon transductions

To ensure the presence of a DNA transposon is a result of active transposition, rather than a passive result of other processes, such as retrotransposition-mediated transductions, we mapped the candidate entries against the known retrotransposons in the ten primate genomes based on their genomic positions. Specifically, the sequences of candidates from the previous step were mapped back onto the host genome using BLAT, followed by removing all entries located within 50 bps to a retrotransposon (excluding entries inserted into a retrotransposon), as such entries could be a result of retrotransposition-mediated transduction. All entries left at this point were considered candidates of “retro-DNAs” for being retrotransposons derived from DNA transposons but apparently using a retrotransposition mechanism.

#### Identification of polyA tails

For each candidate retro-DNA, we retrieved the 10 bp sequence from the 3’ end of the positive-strand (by the DNA transposon consensus sequence). If the sequence contains 6 or more “A”, the entry is considered to have a polyA tail.

### Clustering retro-DNAs to identify unique retro-DNA events

The retro-DNA candidates identified from the last step in the ten primate genomes were subject to a round of “all-against-all” sequence similarity search using BLAT with the sequences of the retro-DNAs plus 100 bp of the flanking region on each side. Entries with 95% or higher sequence similarity across the entirety of the sequences including the flanking sequences were identified as one orthologous cluster, representing one retro-DNA insertion event during the evolution of these primates.

### Estimating the timeline for retro-DNA insertions

A phylogenetic tree of the ten primate genomes plus the marmoset genome as the outgroup was obtained from the TimeTree database (http://www.timetree.org) (Hedges et al. 2006). The treeview program (Page 1996) was used to display the organismal phylogenetic tree. We then added the numbers of non-redundant retro-DNA entries onto the nodes and branches of this tree based on the unique presence of retro-DNAs in the specific genomes or lineages.

### Multiple sequence alignment of retro-DNA and parent sites

We performed multiple sequence alignment for a few selected retro-DNA entries, including their parent sites. We first collected retro-DNA sequences including 100 bp on both flankings, as well as the orthologous sequences of the parent sites from the rest of primate genomes using the online version of MUSCLE (<u>MU</u>ltiple <u>S</u>equence <u>C</u>omparison by <u>L</u>og-<u>E</u>xpectation)(Madeira et al. 2019) from European Bioinformatics Institute website (https://www.ebi.ac.uk/Tools/msa/muscle/).

### Expression analysis of retro-DNAs and their parent copies

RNA-Seq data for the blood and the generic (mixed) samples from chimpanzee, gorilla, crab-eating macaque, rhesus and baboon were retrieved from the Non-Human Primate Reference Transcriptome Resource (NHPRTR)(Pipes et al. 2013) for expression analysis of the retro-DNAs and their parent copies. We also collected data for six human transcriptomes (Shin et al. 2014) and two green monkey transcriptomes (A. J. Jasinska et al. 2013; Anna J. Jasinska et al. 2017). Tophat2 (version 2.1.1) was used to align the RNA-seq reads to the reference primate genomes (D. Kim et al. 2013). Reads mapped to the retro-DNA/parent copies regions were retrieved in fasta format and aligned back to the reference genome using the NCBI BLASTn to ensure that each RNA-seq read was only assigned to only one genomic location based on perfect match, and they were used to calculate the <u>F</u>ragments <u>P</u>er <u>K</u>ilobase of transcript per Million reads (fpkm) values for each DNA transposon entry using an in-house Perl script.

### Data analysis

The data analysis and figure plotting were performed using a combination of Linux shell scripting, R, and Microsoft Excel. The computational analysis was mostly performed on Compute Canada high-performance computing facilities (http://computecanada.ca).

## Results

### Overall profiles of DNA transposons and lineage-specific retro-DNAs in the ten primate genomes

To identify the retro-DNA events in the primate genomes, we first identified the da-DNAs that represent DNA transposons with both the insertion allele and pre-integration allele identifiable in these genomes. These DNA transposons were likely to be the results of relatively recent transposition events, having a low level of sequence divergence and permitting accurate identification of TSDs and TIRs. The starting lists of DNA transposons were based on the RepeatMasker annotation subject to a consolidation process to ensure the accuracy in identifying DNA transposons with both insertion and pre-integration alleles as well as their TSDs. As shown in Table 1, the raw number of DNA transposons in the primate genomes ranged from 392,937 in the marmoset as the lowest to 510,250 in the chimpanzee genome as the highest, averaging at 459,521/genome. After integration, the counts dropped ~18%, leading to less variation in the numbers ranging from 324,288 in marmoset to 421,580 in chimpanzee and averaging at 376,720 DNA transposons per genome. These DNA transposons contributed to a total of ~98 Mbp or ~3.6% of these primate genomes on averages (Table 1). While the numbers and percentages of DNA transposons in these genomes were similar overall with the variation falling within 10% of the average (data not shown), visible differences were also observed based on the integrated DNA entries with marmoset showing the least number and chimpanzee showing the highest (Table 1). Various factors could have impacted the DNA transposons numbers in these genomes, which include, but are not limited to, the differences in the versions of RepeatMasker and the ME reference sequences used for ME annotation, the quality of genome assemblies, and the evolution history of individual genomes.

**Table 1.** Summary of DNA transposons in the 10 primate genomes.

| **Genomes** | **Raw counts** | **integrated counts** | **% count reduction** | **total size (bp)** | **% genome** | **full-length count** | **% full-length** | **diallelic DNA counts** |
| --- | --- | --- | --- | --- | --- | --- | --- | --- |
| **hg38 (human)** | 483,994 | 399,590 | 17 | 102,664,356 | 3.5 | 119,368 | 29.9 | 25,933 |
| **panTro5 (chimp)** | 510,250 | 421,580 | 17 | 107,832,154 | 3.8 | 119,265 | 28.3 | 28,273 |
| **gorGor4 (gorilla)** | 503,480 | 418,454 | 17 | 106,573,049 | 3.8 | 117,263 | 28.0 | 27,386 |
| **ponAbe2 (orangutan)** | 429,467 | 347,471 | 19 | 93,420,030 | 3.4 | 113,425 | 32.6 | 23,923 |
| **nomLeu3 (gibbon)** | 438,800 | 363,738 | 17 | 93,531,426 | 3.6 | 108,334 | 29.8 | 24,206 |
| **macFas5 (crab-eating macaque)** | 443,909 | 359,802 | 19 | 94,910,440 | 3.5 | 109,444 | 30.4 | 26,218 |
| **rheMac8 (rhesus)** | 486,991 | 401,546 | 18 | 102,546,356 | 3.7 | 111,558 | 27.8 | 28,149 |
| **papAnu2 (baboon)** | 459,662 | 369,684 | 20 | 97,943,467 | 3.7 | 109,523 | 29.6 | 25,844 |
| **chlSab2 (green monkey)** | 445,724 | 361,048 | 19 | 95,097,218 | 3.5 | 108,139 | 30.0 | 26,252 |
| **calJac3 (marmoset)** | 392,937 | 324,288 | 17 | 83,220,943 | 3.2 | 91,946 | 28.4 | 34,901 |
| **Average** | 459,521 | 376,720 | 18 | 97,773,944 | 3.6 | 110,827 | 29.5 | 27,109 |
| **\*: full-length is defined as >=90% of consensus** |  |  |  |  |  |  |  |  |

Using a multi-way comparative genomics approach modified from our previous analysis of human-specific MEs (Tang et al. 2018), we identified a total of 271,085 da-DNAs in the ten primate genomes (Table 1). Specifically, for each da-DNA, we require the presence of a preintegration allele in at least one of the nine remaining genomes. As shown in Table 1, the number of da-DNAs varies from 23,923 in the orangutan genome as the lowest to 34,901 in the marmoset as the highest, averaging at 27,109 for the ten genomes. The largest number of da-DNAs in the marmoset is expected as marmoset has the largest evolutionary distance from the remaining primate species. Notable differences were also seen between genomes with mutually closest evolutionary relationship in the group, making the numbers directly comparable for the paired genomes. For example, between the human and chimpanzee pair, the chimpanzee genome has more than 10% of da-DNAs than the human genome (28,273 vs. 25,933), while between the two macaques, the rhesus genome has ~10% more than the crab-eating macaque genome (28,149 vs. 26,218) (Table 1). Interestingly, this difference is much less than that for the species-specific non-LTRs, which shows crab-eating macaque genome having a much lower retrotransposition activity than the rhesus genome (Tang and Liang 2019). This may indicate that majority of these da-DNAs were generated by a mechanism different from retrotransposition.

By the composition in DNA transposon type, majority of the da-DNAs belong to the hAT and TcMar superfamilies (Table S1, Fig. S1). The two hAT families, hAT-Charlie and hAT-Tip100, contributed to ~57% of da-DNAs in all genomes with the hAT-Charlie family alone contributing to ~50% of all da-DNAs. The two TcMar families, TcMar-Tigger and TcMar-Mariner, contribute ~30% of da-DNAs, while the remaining families contributed to ~10% of da-DNAs. This composition pattern was quite similar among all genomes, except for the orangutan genome, which has less da-DNAs from the TcMar-Trigger and the hAT-Tip100 families but more from families other than the hAT and TcMar superfamilies (Fig. S1).

### Retro-DNAs in the primate genomes show non-LTR retrotransposon sequence characteristics

While analyzing these da-DNAs in detail for understanding the possible mechanisms involved, we came across an unusual case of DNA transposon located at *chr4:146335052-146335253* of the human genome (GRCh38) as being a human-specific ME (Fig. 1A). The 201 bp DNA transposon fragment was annotated by RepeatMasker as a *Tigger7* element from the *TcMar-Tigger* family. As shown in the multiple sequence alignments (Fig. S2) with its orthologous sequences from other eight primate genomes (not identifiable in marmoset genome), the *Tigger7* element was absent from the orthologous sites of all non-human primate genomes, confirming it as an authentic case of human-specific ME. Interestingly, this insertion has a 15 bp TSD “AAGAGTCCTGGATCC/AAGAGTCCTGGATCA” that was much longer than TSDs for DNA transposons, and it has no identifiable TIR typical of a DNA transposon (Fig. 1A). Furthermore, it has a 27 bp polyA tail at the 3’-end of the insertion sequence and a predicted polyadenylation signal “ATTAAA” before the polyA tail. Despite being part of a *Tigger7* DNA transposon sequence, all these features point this insertion to be a non-LTR retrotransposon rather than a canonical *Tigger7* DNA transposon, which is expected to have TIRs and 2 bp (TA) TSDs. Because it is a DNA transposon sequence with the characteristics of a non-LTR retrotransposon, something not reported before, we named it as a retro-DNA for being a retrotransposon-like element derived from a DNA transposon sequence.

**Figure 1.**
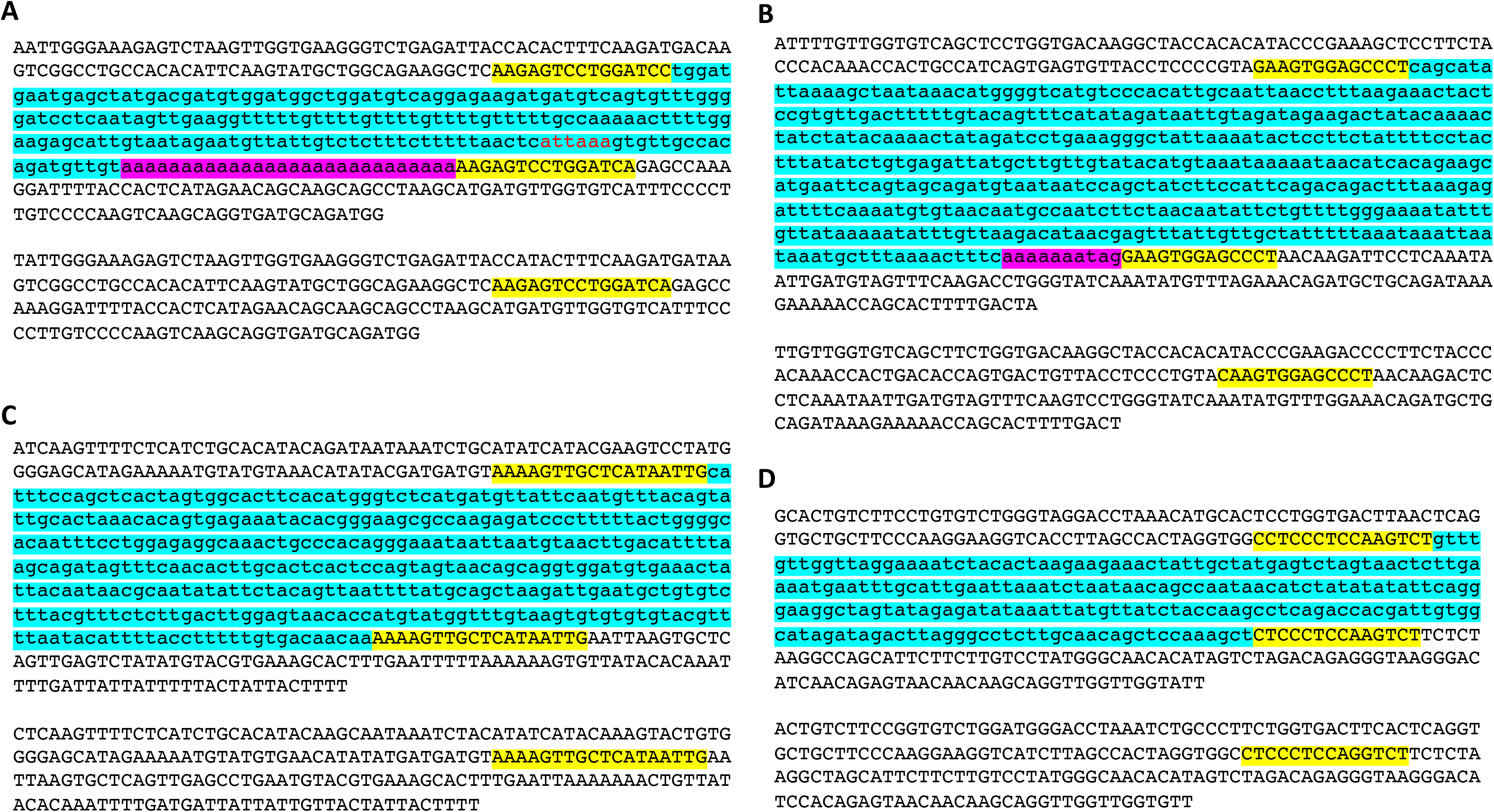
Examples of retro-DNAs in different primate genomes. A. A retro-DNA from the human genome (hg38_chr4:146335052-146335253) with the pre-integration allele from the chimpanzee genome (panTro5_chr4:38758218-38758438). B. A retro-DNA from the green monkey genome (chlSab2_chr8:30005081-30005527) with the pre-integration allele from the gibbon genome (nomLeu3_chr8:37535028-37535236); C. A retro-DNA located from the green monkey genome (chlSab2_chrX:73456937-73457324) with the pre-integration allele from the orangutan genome (ponAbe2_chrX:82896142-82896360). D. A retro-DNA located from the human genome (hg38_chr4:38758216-38758442) with the pre-integration allele from green monkey genome (chlSab2_chr27:11529606-11529817). In each panel, the sequence at the top is the insertion allele containing the retro-DNA, and the sequence at the bottom is the preintegration allele without the retro-DNA. The yellow boxes indicate TSDs, the blue boxes indicate the DNA transposon sequences, while the purple boxes indicate possible polyA tail sequences.

Following discovering this case of retro-DNA, we searched the human genome and other genomes and identified more similar cases, as exampled in Fig. 1B-D. For instance, a 446 bp *Charlie1a* fragment from the *hAT-Charlie* family was identified as a retro-DNA in three primate genomes (the green monkey, rhesus, and crab-eating macaque genomes with the locations being chlSab2_chr8:30005081-30005527, rheMac8_chr8:31992158-31992606, and macFas5_chr8:32527581-32528029, respectively). This entry has a 13 bp TSD “GAAGTGGAGCCCT” and no TIRs (Fig. 1B). As shown in Fig. S3, the 446 bp *Charlie1a* fragment was absent in the orthologous regions of the remaining seven primate genomes, which could be explained as a lineage-specific insertion event that happened in the last common ancestor of green monkey, rhesus, and crab-eating macaque. Also, it appears that this retro-DNA sequence in these three genomes had been subject to variations in the polyA tails as having different lengths, indicating its relatively older age as a lineage-specific da-DNA in comparison to the species-specific element as exampled in Fig. S2.

By requiring the presence of longer TSDs (≥8 bp) and the absence of TIRs, we identified a total of 1,750 retro-DNA entries from the da-DNAs using a workflow shown in Fig. 2. By classification, these retro-DNAs consist of 847, 478, 156, 74, and 195 entries from the hAT-Charlie, TcMar-Tigger, hAT-Tip100, TcMar-Mariner, and other families, respectively. As seen in Table 2, these 1,750 retro-DNA entries cover all ten genomes and can be clustered into 748 unique retro-DNA insertion events. It is worth noting that our list of retro-DNAs may suffer a certain level of false-negatives and false-positives due to the uses of a set of criteria which may not be very optimal and due to challenges associated with the analysis of transposable elements as well as the deficiencies of the resources. The latter includes the quality of the reference genomes and the RepeatMasker annotation, especially for the non-human primates as discussed in our recent study (Tang and Liang 2019).

**Figure 2.**
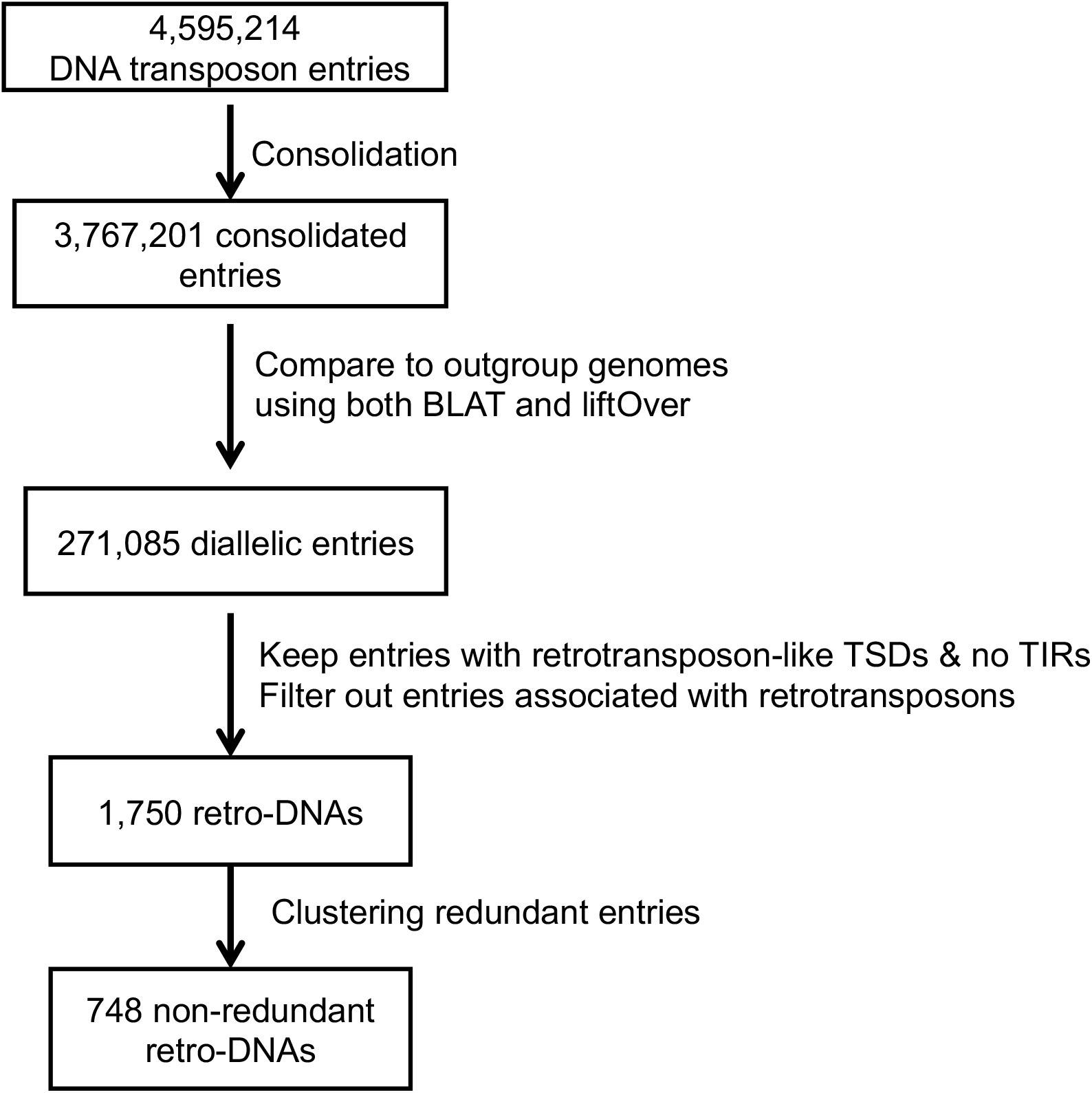
Flow chart for identification of retro-DNAs.

**Table 2.** Retro-DNAs in the 10 primate genomes.

| **Table 2. Retro-DNAs in the 10 primate genomes** |  |  |  |  |  |  |  |  |  |  |  |  |
| --- | --- | --- | --- | --- | --- | --- | --- | --- | --- | --- | --- | --- |
| **DNA transposon family** | **Human** | **Chimpanzee** | **Gorilla** | **Orangutan** | **Gibbon** | **Crab-eating macaque** | **Rhesus** | **Baboon** | **Green monkey** | **Marmoset** | **Total** | **Total (nr)** |
| hAT-Charlie | 100 | 108 | 99 | 58 | 72 | 76 | 79 | 76 | 78 | 101 | 847 | 317 |
| hAT-Tip100 | 19 | 17 | 18 | 10 | 19 | 16 | 16 | 13 | 13 | 15 | 156 | 63 |
| TcMar-Tigger | 44 | 51 | 49 | 28 | 47 | 49 | 57 | 36 | 58 | 59 | 478 | 221 |
| TcMar-Mariner | 7 | 8 | 8 | 2 | 6 | 7 | 8 | 7 | 6 | 15 | 74 | 34 |
| Other | 17 | 18 | 17 | 56 | 12 | 15 | 17 | 11 | 15 | 17 | 195 | 113 |
| **All Retro-DNAs** | **187** | **202** | **191** | **154** | **156** | **163** | **177** | **143** | **170** | **207** | **1,750** | **748** |

By sequence length, these 748 retro-DNA entries averaged at 209 bp (±190 bp) and represented only part of DNA transposons in all cases, covering ~21% of their consensus sequences (Table 3). While the consensus sequences for DNA transposon families differ in length significantly from 380 bp for TcMar-Mariner to 1506 bp for hAT-Tip100, the average length of retro-DNAs seems to be relatively consistent among the families, ranging from 122 bp for TcMar-Mariner to 251 bp for TcMar-Tigger. Nevertheless, in general, the retro-DNAs from the longer families do have a longer average length than those from the shorter families, despite those from families with longer consensus sequences (e.g. hAT-Tip100) having lower proportion of their consensus sequences than those with shorter consensus sequences (e.g. TcMar-Mariner)(Table 3).

**Table 3.**
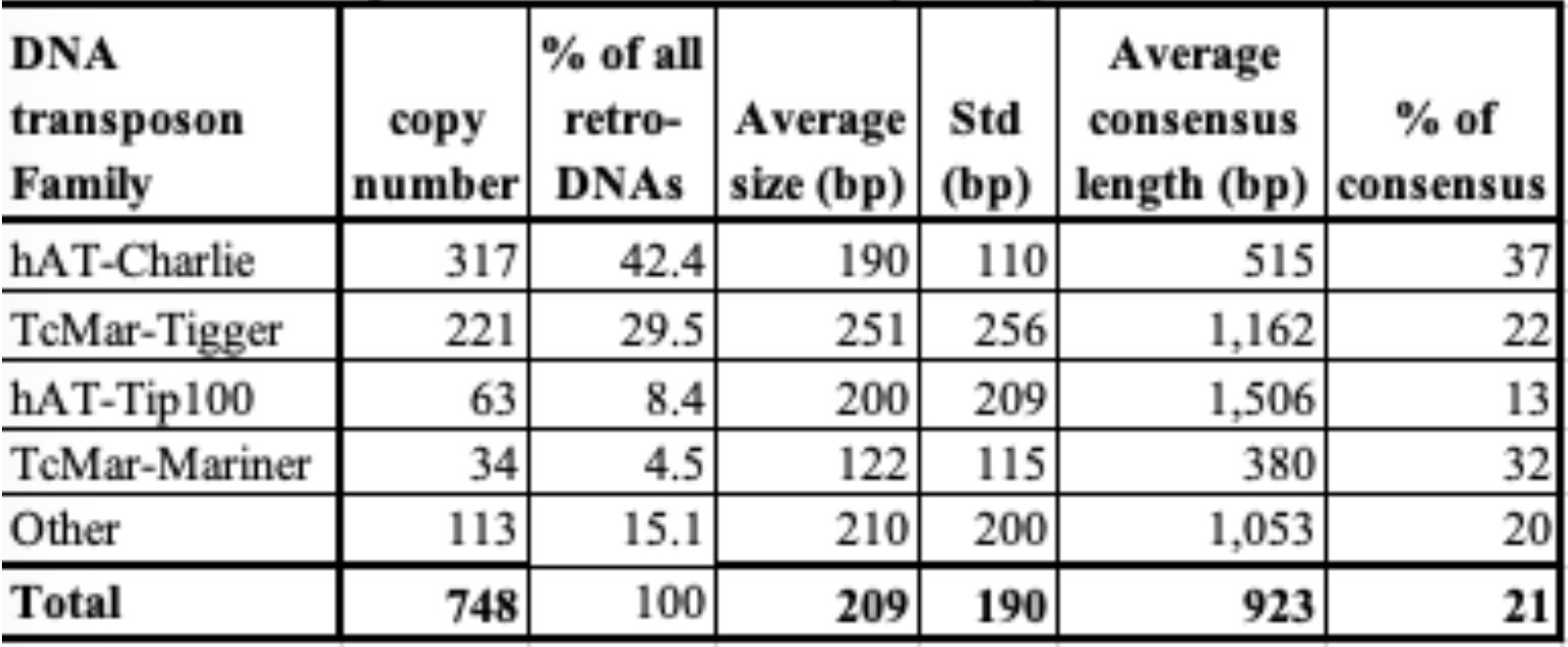
The composition of retro-DNA by family and the size information.

| **Table 3. The composition of retro-DNAs by family and the size information** |  |  |  |  |  |  |
| --- | --- | --- | --- | --- | --- | --- |
| **DNA transposon Family** | **copy number** | **% of all retro-DNAs** | **Average size (bp)** | **Std (bp)** | **Average consensus length (bp)** | **% of consensus** |
| hAT-Charlie | 317 | 42.4 | 190 | 110 | 515 | 37 |
| TcMar-Tigger | 221 | 29.5 | 251 | 256 | 1,162 | 22 |
| hAT-Tip100 | 63 | 8.4 | 200 | 209 | 1,506 | 13 |
| TcMar-Mariner | 34 | 4.5 | 122 | 115 | 380 | 32 |
| Other | 113 | 15.1 | 210 | 200 | 1,053 | 20 |
| **Total** | **748** | **100** | **209** | **190** | **923** | **21** |

Additionally, we examined whether there are any hotspots on the consensus sequences that are used as the source sequences of these retro-DNAs. By using the retro-DNA entries from the Tigger1 DNA transposon subfamily, the largest subfamily containing 41 non-redundant retro-DNAs, we generated a frequency plot showing the usage of the consensus sequences by the retro-DNAs. As illustrated in Fig. S4, while all regions of the consensus sequence were used by the 41 retro-DNAs, the frequency ranges from 2.4% to 29.3%, showing that a few regions including those from ~1310-1440 bp and ~1840-2240 bp of consensus sequence had been used more frequent than the rest of the regions.

As shown in Table S2, from the total 748 retro-DNAs we identified a total of 176 non-redundant retro-DNA entries carrying a potential polyA tail. We speculate that the relatively low percentage (23.5%) of entries with a polyA tail might be partially due to post-insertion mutations in the polyA sequences, which are more prone to random mutations than other regions. The complete list of the 748 non-redundant retro-DNA entries with their genomic coordinates in all applicable genomes was provided in Supplementary file S1. For these retro-DNA insertion events, we further examined the sequence motifs at the insertion sites and the TSD length distribution pattern. As shown in Fig. 3A, a sequence motif of ‘TT/AAAA’, which was the same as the motif for *Alu*s, L1s, and SVAs (Goodier 2016; Tang et al. 2018; Wang et al. 2006), was clearly seen despite the signal being much weaker. This, nevertheless, is a strong indication of their use of the L1-based non-LTR retrotransposition TPRT mechanism (Cost and Boeke 1998; Jurka 1997). As a further support, the TSD length distribution peaks at 8 bp. Despite being shorter than the peak around 15 bp, the major peak observed for non-LTR retrotransposons, this 8 bp peak is similar to a secondary peak for the TSD lengths of human specific L1s (Tang et al. 2018).

### The patterns of retro-DNAs and their parent sites in genome distribution and expression

To assess the potential functional impact of these retro-DNAs, we examined their gene context based on the Ensemble gene annotation for these genomes (Release 95 for all genomes except Release 90 and 91 were for baboon and marmoset, respectively)(Zerbino et al. 2018). A total of 698 retro-DNAs, representing ~40% of the 1,750 retro-DNAs, are located within different genic regions, including non-coding RNAs, intron regions, untranslated regions, and promoter regions for 734 transcripts representing 414 unique genes (Table 4 & Table S3). The majority of these retro-DNAs were located within the intron regions (699/734), while 27 entries were inserted into promoter regions and the untranslated regions. The presence of these retro-DNAs in the genic regions provides the potential for them to impact gene regulation or splicing.

**Table 4.** The numbers of retro-DNAs located in the genic regions in the 10 primate genomes.

| **Table 4. The numbers of retro-DNAs located in the genic regions in the ten primate genomes** |  |  |  |  |  |  |  |  |  |  |  |
| --- | --- | --- | --- | --- | --- | --- | --- | --- | --- | --- | --- |
| **Genic region\*** | **Human** | **Chimpanzee** | **Gorilla** | **Orangutan** | **Gibbon** | **Crab-eating macaque** | **Rhesus** | **Baboon** | **Green monkey** | **Marmoset** | **Total** |
| NR | 4 | 1 | 1 | 1 | 0 | 0 | 1 | 0 | 0 | 0 | 8 |
| Promoter | 9 | 5 | 2 | 1 | 0 | 1 | 1 | 0 | 0 | 3 | 22 |
| 5' UTR | 0 | 1 | 0 | 0 | 0 | 1 | 0 | 0 | 0 | 0 | 2 |
| 3' UTR | 1 | 0 | 0 | 0 | 0 | 0 | 0 | 0 | 0 | 2 | 3 |
| Intron | 114 | 78 | 70 | 60 | 61 | 62 | 67 | 42 | 53 | 92 | 699 |
| **Total** | **128** | **85** | **73** | **62** | **61** | **64** | **69** | **42** | **53** | **97** | **734** |
| **Total (nr)** | **109** | **82** | **70** | **60** | **61** | **63** | **67** | **42** | **53** | **91** | **698** |

\*, NR: non-coding RNA; UTR: untranslated region

We also examined the timeline of these retro-DNA insertion events by mapping them onto a phylogenetic tree of these primates based on the data in the TimeTree database (http://www.timetree.org)(Hedges et al. 2006). As shown in Fig. 4A (insert), 450 or 60.2% of these retro-DNAs appeared to be species-specific by being uniquely present in only one genome, while another 295 were found in multiple genomes as being lineage-specific. On average a retro-DNA is shared among two genomes, suggesting an average age older than the species-specific MEs reported in our earlier study (Tang and Liang 2019). Some manual corrections were made for placing the lineage-specific retro-DNAs on the phylogenetic tree. For example, 7 retro-DNAs are found to be shared between human and gorilla but not in chimpanzee, and we decided to include these entries on the branch common for these three genomes. We argue that this manual correction is necessary as retro-DNA identification can suffer false negatives from the insufficient sequence assembly quality for the non-human primate genomes. As shown in Fig. 4B, the number of retro-DNA insertion events seems to show a positive linear correlation with the relative evolutionary spans of the species and lineages (R^2^ = 0.5463), suggesting that these retro-DNA insertion events have occurred at a low but relatively consistent rate during primate evolution.

**Figure 3.**
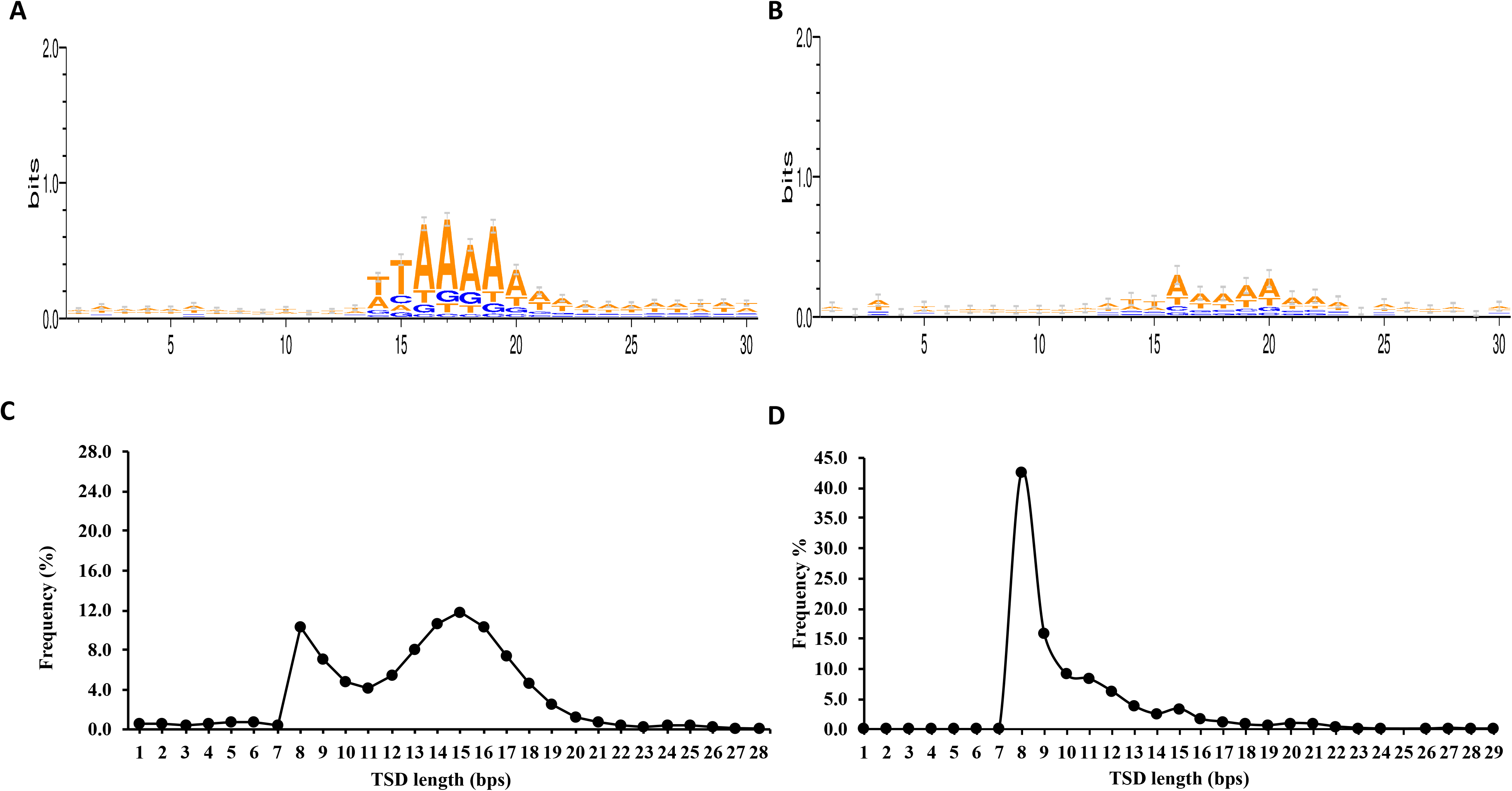
Sequence motifs of pre-integration sites and TSD length distribution pattern for retro-DNAs. A. Sequence motif logos for human-specific L1s at the integration sites, adopted from authors’ publication (Tang et al, 2018). B. Sequence motif logos for retro-DNAs at the integration sites. C. A line plot showing the distribution of TSD length for human-specific L1s, adopted from authors’ publication (Tang et al, 2018). D. A line plot showing the distribution of TSD length for retro-DNAs.

Further, we identified the potential parent sites for these retro-DNA entries by performing sequence similarity search using these retro-DNAs as query sequences against each primate genome. For each retro-DNA, the best non-self-match was selected as its potential parent site. As shown in Table S4, we have identified a total of 715 potential parent sites for the 1,750 retro-DNA entries. This converts to 325 non-redundant entries of the 748 unique retro-DNAs. The failure in finding the parent copies for the remaining entries could be due to the loss of the parent copy as result of genomic rearrangements or due to incomplete coverage of the genome sequences. As for retro-DNAs, we have examined the gene context for these potential parent sites. As shown in Table S5, 351 (49.1%) of these redundant potential retro-DNA parent sites were located within 410 different genic regions including 13 coding sequences, 19 non-coding RNAs, 112 promoter regions, one 5’ UTR, four 3’UTRs and 274 intron regions, which collectively represent 371 unique genes/transcripts. In these cases, the transcripts of these potential parent sites, likely as part of the transcripts or splicing products of their host genes might have had the chance to hijack the L1 TPRT machinery as in the case of processed pseudogenes to generate the retro-DNAs. The ratio of genic entries (49.1%) is higher for parent sites than that for retro-DNAs (~40%), and its implication is discussed in later sections.

We also examined the expression level of retro-DNAs and their potential parent sites using RNA-seq data from <u>N</u>on-<u>H</u>uman <u>P</u>rimate <u>R</u>eference <u>TR</u>anscriptome (NHPRTR) dataset (Pipes et al. 2013) and two other studies (Anna J. Jasinska et al. 2017; Shin et al. 2014) to see if any of these entries have any transcriptional activities in the current primate genomes. For this, we collected a total of 21 transcriptomes for seven primates, excluding orangutan, gibbon, and marmoset, for which no transcriptome data is available. To minimize the false positives due to the high sequence similarity among members in the same family, we included only the reads with the perfect match to the retro-DNAs or their parent site regions in the primate genomes and with each read used only once in calculating the expression level. However, since the specific transcriptome sequences can diverge from the corresponding reference genomes due to intraspecies variations, we believe this process has inevitably introduced false negatives in the results and therefore lead to an underestimation of the retro-DNAs and parent sites’ expression level. As seen in Table 5 & S6, 966 loci from the 1,750 retro-DNA and 715 parent sites in these seven primate genomes were shown to have a certain level of expression ranging in fpkm value from 0.0003 to 27.30.

**Table 5.** The numbers of expressed retro-DNAs and parent sites in 21 primate transcriptomes.

| Species | # of RNA-seq sets | retro-DNAs |  |  | parent sites |  |  |
| --- | --- | --- | --- | --- | --- | --- | --- |
|  |  | entry # | # of expressed | % | entry # | # of expressed | % |
| Human | 6 | 187 | 93 | 49.7 | 98 | 57 | 58.2 |
| Chimpanzee | 2 | 202 | 99 | 49.0 | 101 | 67 | 66.3 |
| Gorilla | 1 | 191 | 55 | 28.8 | 99 | 42 | 42.4 |
| Rhesus | 4 | 177 | 97 | 54.8 | 64 | 46 | 71.9 |
| Crab-eating macaque | 4 | 163 | 115 | 70.6 | 63 | 55 | 87.3 |
| Baboon | 2 | 143 | 68 | 47.6 | 53 | 34 | 64.2 |
| Green monkey | 2 | 170 | 90 | 52.9 | 62 | 48 | 77.4 |
| Total | 19 | 1063 | 527 | 49.6 | 478 | 301 | 63.0 |

We further investigated the relationship between retro-DNAs and their parent sites based on their expression levels. Specifically, three human testis transcriptome samples (SRR2040581, SRR2040582, SRR2040583) retrieved from the NCBI SRA (<u>S</u>equence <u>R</u>ead <u>A</u>rchive; https://www.ncbi.nlm.nih.gov/sra) were used for analyzing expression level of the retro-DNA/parent site pairs. As shown in Fig. 5A, a total of 66 retro-DNA/parent site pairs were shown to have a certain level of expression (fpkm > 0) for either the retro-DNA or the parent site among the three human testis samples. Notably, within these 66 retro-DNA/parent site pairs, 57 parent sites showing expressed (fpkm > 0) compared to only 42 entries for retro-DNAs (Table S4 & S6, Fig. 5A). This might be due to the fact that the generation of a retro-DNA requires the expression of its parent site, while a retro-DNA itself may not be expressive depending on its landing location, which is random. Therefore, a higher ratio of transcriptionally active sites can be expected for the parent sites than the progeny (retro-DNA) sites. More interestingly, we noticed that the two parent sites responsible for multiple retro-DNA entries were shown to have the highest levels of expression among the parent sites (Fig. 5A, labeled in brackets). This may suggest that the expression level of the parent sites is positively correlated to their potential in generating retro-DNAs and that there seems to be no relationship between the expression levels of the parent sites and the progeny retro-DNA sites. Furthermore, the ongoing expression of the parent sites suggests that they can potentially generate more retro-DNAs in the future.

We also examined and compared the expression levels of retro-DNAs and their parent sites in the three human testis transcriptomes by classification in gene contest. As shown in Fig. 5B, the average fpkm values of the parent sites are always higher than the average fpkm values of the retro-DNA entries as a whole group or divided into genic and intergenic regions. In addition, the entries located within genic regions showed higher expression than the ones located outside the genic regions for both retro-DNAs and the parent sites (Fig. 5B), suggesting that entries located in the genic region may have more opportunities to be expressed passively as part of the host gene expression. This difference is larger for retro-DNAs than for the parent sites, likely because parent sites have to be expressed regardless of their position in order to be able to generate new copies and their expression might be driven by other factors if located outside genes. None of these differences are statistically significant, likely due to the small sample size.

## Discussions

### Retro-DNAs as a new type of retrotransposons derived from DNA transposons

In this study, we focused our attention on a small number of species-specific DNA transposons identified in primate genomes using our computational comparative genomics pipelines which revealed unprecedented numbers of species-specific retrotransposons in the human genome and seven other genomes (Tang et al. 2018; Tang and Liang 2019). Unlike for the retrotransposons, for which the ongoing activity during primate evolution and in the current genomes have been well established (Goodier 2016; Jordan et al. 2018; Tang and Liang 2019), the presence of species-specific DNA transposons in these primate genomes presents a puzzle. It cannot be answered by existing literature, because DNA transposons are thought to have become inactive about 37 Mya ago (Feschotte and Pritham 2007; Pace Ii and Feschotte 2007), meaning that no canonical DNA transposition activity could have existed during the evolution of these primate genomes. In trying to understand the mechanism underlying these mystery species-specific DNA transposon insertions, we examined the sequence features manually and spotted a few interesting entries as exemplified by the case shown in Fig. 1A, which shows clear characteristics of non-LTR retrotransposons by having longer TSDs and presence of a polyA tail, while lacking TIRs that are the hallmark of new DNA transposon insertions. The remaining cases shown in Fig. 1 have the same non-LTR features, but not necessarily have the typical polyA tail. For their non-LTR retrotransposon characteristics, we name them as “retro-DNA” as retrotransposons derived from DNA transposons. We then performed systematic analysis to look for more of such “retro-DNA” cases.

For this, we expanded from the strict species-specific MEs, which are defined as those present in only one of the primate genomes (Tang et al. 2018; Tang and Liang 2019), to diallelic DNA transposons or da-DNAs, which are defined as those with a pre-integration site (i.e., the orthologous region without the DNA transposon) present in at least one of the ten genomes. We obtained a total of 271,085 da-DNAs, and from these we then specifically searched for retro-DNA cases which have long TSDs (>=8bp) and the absences of the TIRs using a protocol shown in Fig. 2. This led to the identification of 1,750 of retro-DNA cases, which represent 748 unique events, covering all ten primate genomes with the over half being species-specific and the other half being lineage-specific at different levels on the evolution tree (Fig. 4A). Our results indicate that the presence of retro-DNAs is not limited to the human genome but can be found in all ten primate genomes included in this study and along different evolution stages of these primates. Furthermore, these retro-DNAs are not limited to one DNA transposon family but cover all major DNA transposon families, suggesting that the existence of such “retro-DNAs”, a novel type of retrotransposons, is not just limited to incidental rare cases, but is rather the product of a consistent mechanism existed in all these primate genomes.

**Figure 4.**
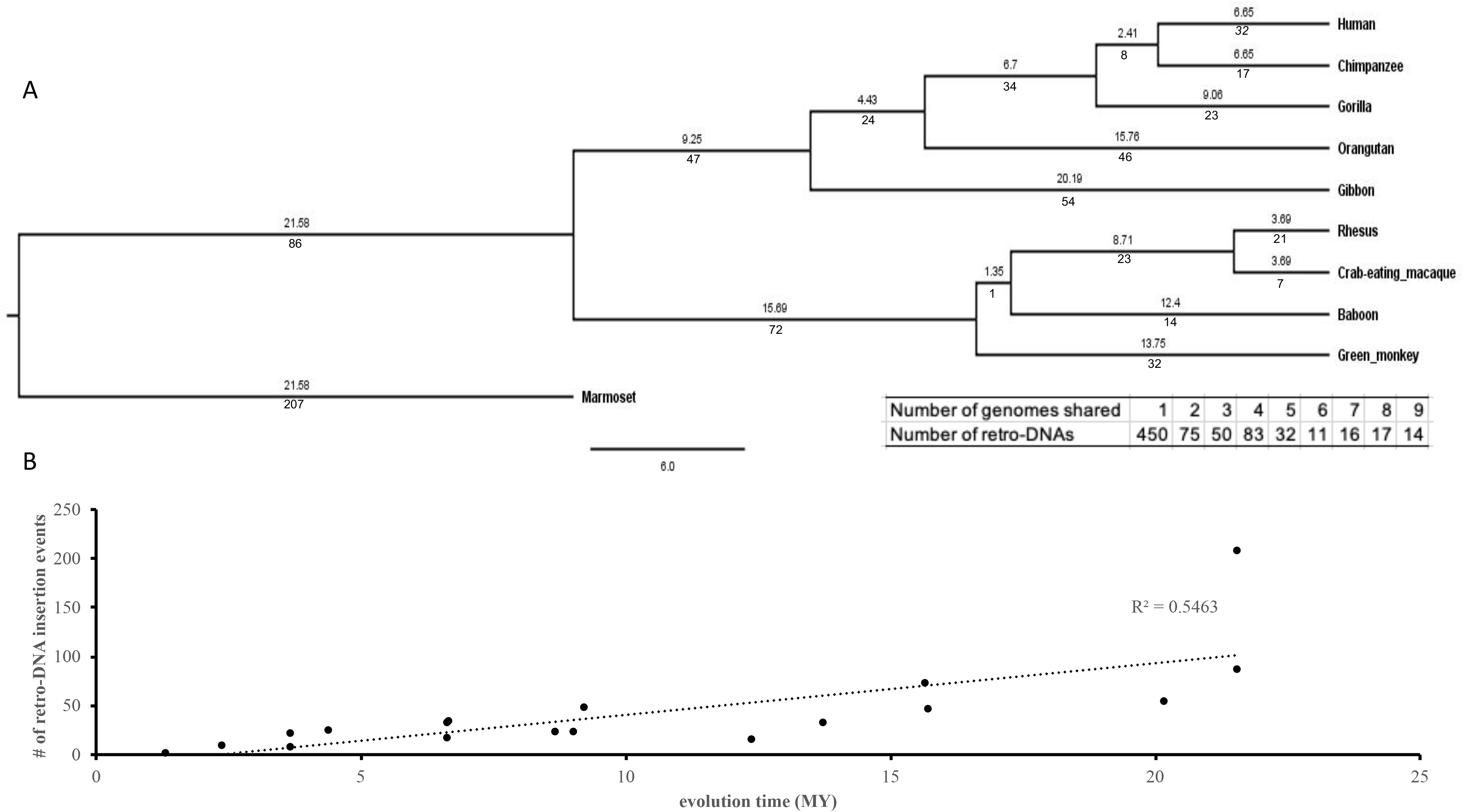
The timeline for the generation of retro-DNAs during the evolution of the ten primate genomes. A. A rooted phylogenetic tree of the ten primate genomes from the TimeTree database(http://www.timetree.org/). The numeric values below each branch represent the number of retro-DNA insertion events happened during the corresponding period of primate evolution. The numeric value above each branch represents the millions of years (Mya) for that branch. B. A scatter plot between the number of retro-DNA insertion events and the evolutionary time based on the data in panel A. The trend line shows that the number of retro-DNA insertion events is positively correlated with the relative evolutionary distance with R^2^ = 0.5463.

### The likely mechanism underlying the generation of retro-DNAs

Several lines of evidence from our results guided us to propose that these retro-DNAs were the products of the L1-based TPRT machinery, similar as for the known non-autonomous non-LTR retrotransposons, i.e., SINEs, SVAs and processed pseudogenes (Cost and Boeke 1998; Jurka 1997; Mita and Boeke 2016; Tang et al. 2018; Xing et al. 2006). The major pieces of evidence include the presence of the “TT/AAAA” sequence motif at the insertion sites and the long TSDs. As seen in Fig. 3A, the integration sites for the 748 retro-DNAs display a core sequence motif of “TT/AAAA”, which is identical to the insertion site sequence motif observed for non-LTR retrotranposons in the human genome (Jurka 1997; Tang et al. 2018; Wang et al. 2006). The length distribution of the TSDs for these retro-DNAs, as shown in Fig. 3B, shows a dominant peak at 8bp, which is much longer than that of TSDs typically found for DNA transposons (2 bp) and is similar to the secondary peak of TSD length observed for the human-specific L1s (Tang et al. 2018).

As additional pieces of evidence supporting our proposal, the presence of parent sites in the same genome for a high proportion of the retro-DNAs (325/748 or 43.5%) indicates their use of a “copy-and-paste” rather than the “cut-and-paste” mechanism used by canonical DNA transposons. Furthermore, the presence of a polyA tail in many (176/748 or 23.5%) of these retro-DNAs provides additonal support for the use of L1-based TPRT mechanism.

It is worth noting that, as described above, while there is sufficient similarity in sequence features between these retro-DNAs and the known non-LTR retrotranspsons for treating these retro-DNAs as a new type of non-LTR retrotranspons, unique aspects of these retro-DNAs are also visable. These include the missing of the major TSD length peak at 15 bp observed for non-LTR retrotransposons, the low percentage of entries with a polyA tail, and the weaker signal of the sequence motif, “TT/AAAA”, at the integration sites. All of these characteristics might be contributed by the relative older average age of these retro-DNAs as shown by the relatively high percentage (298/748 or ~40%) of being lineage-specific (Fig. 1A) than the non-LTR retrotransposons used in most prior studies for analysis of the non-LTR integration site sequence motifs (Cost and Boeke 1998; Jurka 1997; Mita and Boeke 2016; Tang et al. 2018; Xing et al. 2006). In other words, the older age of the retro-DNAs leads to higher sequence divergence, which in turn lowers the sensitivity for detecting all of these sequence features. An additional reason for the weaker signal in the insertion site sequence motif for the retro-DNAs could be due to the small sample size. In the meantime, it is also possible that these characteristics may suggest that some differences may exist in the detailed retrotransposition process of these DNA transposons, likely the interaction between the retro-DNA transcripts and the ORF1 and ORF2 proteins.

### The relative retro-DNA activity during primate evolution

In comparison with the other types of non-autonomous non-LTR retrotransposons, including Alus, SVAs, and processed pseudogenes, in the primates (Bennett et al. 2008; Goodier 2016; Lander et al. 2001; Tang and Liang 2019), the number of retro-DNAs per genome is much lower, averaging below 200 per genome (Table S2). This number is even much lower than that of processed pseudogenes, which repesent the smallest class of non-LTR retrotransposons with 10,190 in the human genome (Tutar 2012). We reason that the small copy numbers of retro-DNAs may be mainly attributing to one factor, which is the lack of internal promoters to drive their own transcription, leading to an overall low level of their transcripts available for retrotransposition. This is in agreement with the fact that there is no clear hotspot in the DNA transposon consensus sequences used in generating retro-DNAs as shown in Fig. S4 for Tigger 1. Should there be internal promoters driving the transcription, we would expect to observe one or more clear dominant peaks in the frequence of the regions used for retro-DNAs.

Without the ability in driving their own transcription, the only way for DNA transposons to get transcribed is to be become part of genes and get transcribed as a part of the host gene transcripts. If this is how retro-DNAs were generated, then we would expect to see a high percentage of retro-DNAs having their parent sites located in the genic regions, more specifically in the transcribed regions, i.e. exons and introns. By examining the gene context, 351 of the 715 parent sites (49.0%) for the retro-DNAs locate in 371 unique genes/transcripts in the ten primate genomes. This ratio is higher than the ratio of all DNA transposons in the genic regions (39%, detailed data not shown), thus supporting the role of passive expression in generating these retro-DNAs.

Following the same rationale, we would expect that on average the parent sites should have a higher expression level than that of retro-DNAs since the parent sites were selected to be biased for those in the genic regions, while the location of the retro-DNAs is more or less random, leading to the latter having a relative lower proportion in genic regions than the parent sites as observed (40% vs. 49%) (Table 4, Table S5). The observed expresion pattern seems to support our prediction, since among the 66 retro-DNA/parent site pairs, 57 pairs have parent sites with fpkm > 0 compared to only 42 expressed entries for retro-DNAs (Fig. 5A). Additionally, we identified two parent sites, which are potentially responsible for generating multiple retro-DNA entries, and they showed the highest levels of expression among the parent sites (Fig. 5A). Among these two parent sites, the one potentially responsible for two copies of retro-DNAs showed higher expression than that responsible for two retro-DNAs (Fig. 5A). By comparing the expression levels of all parent sites with that of retro-DNAs in the human genome, we can see an overall higher expression for the parent sites (Fig. 5B), and this is also true when comparing the two groups of sites in the genic and intergenic regions (Fig. 5B). Furthermore, the expression level of parent sites in the genic regions is much higher than their counterparts in the intergenic regions as expected (Fig. 5B).

**Figure 5.**
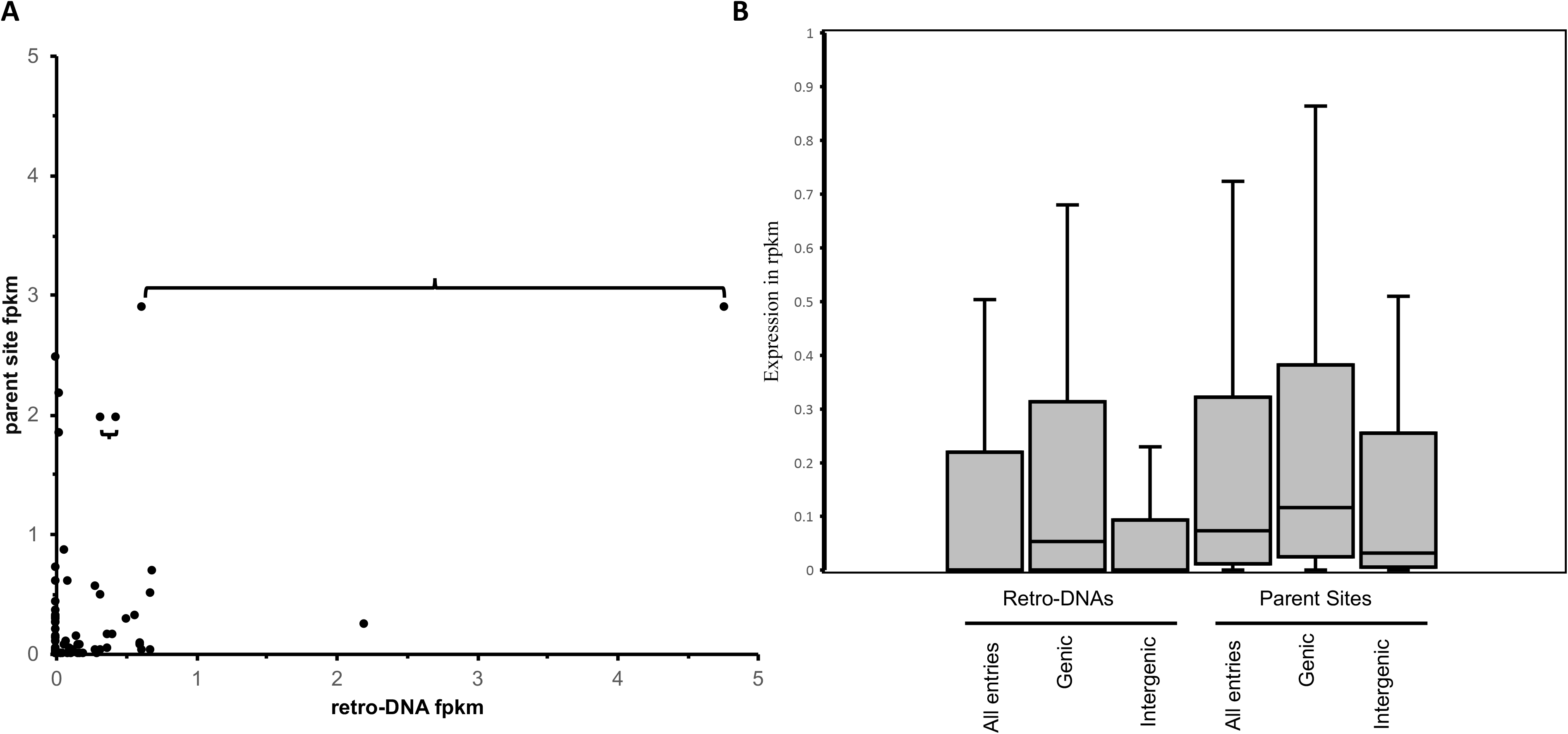
The expression (fpkm) of retro-DNAs and their parent sites in three human testis transcriptomes. A. A scatter plot based on 66 retro-DNA/parent site pairs which show a certain level of expression (fpkm > 0) for the retro-DNA and/or parent site. The retro-DNAs connected by brackets indicate are entries possibly from the same parent copy. B. Box plots showing the expression levels of the 66 retro-DNAs and parent sites divided into genic and intergenic groups. Expression data was based on the average fpkm value in the three human testis transcriptomes.

The use of ten primate genomes, representing several lineages spanning a certain time span in primate evolution, allowed us to examine whether there is any positive correlation between the length of evolutionary span and the number of retro-DNA insertion events. As shown in Fig. 4B, a positive correlation between the two (R^2^ = 0.5463) is observed, suggesting that the generation of retro-DNAs is relatively steady during the evolution of this group of primates. Furthermore, the fact that many of the retro-DNA parent sites, as well as 966 of the 1773 (~54.5%) retro-DNAs show certain levels of expression in the seven primate transcriptomes (Table 5 & S6) suggests the possible ongoing activity of retro-DNA generation from the parent sites and retro-DNAs.

### The functional potential of retro-DNAs

As shown in Table 4 & S3, 698 retro-DNA entries (redundant across genomes), representing ~40% of the 1,750 retro-DNAs, are located within 734 different genic regions, including non-coding RNAs, introns, untranslated regions, and promoter regions for 414 unique genes in the ten primate genomes. Furthermore, 8 entries of these retro-DNAs contribute to part of transcripts, despite none found to be in CDS regions. Therefore, we can speculate that these retro-DNAs may have potential impact on gene function via the regulation of transcription and/or splicing.

### Conclusions and future perspectives

In this study, through a comparative analysis of ten primate genomes including the human genome, we identified a new type of non-autonomous non-LTR retrotransposons derived from DNA transposon sequences. Named as “retro-DNAs”, these elements represent the 5^th^ type of non-LTR retrotransposons after LINE, SINE, SVA, and processed pseudogene, very likely using the same L1-based TPRT mechanism. The generation of these retro-DNAs serves to propagate DNA transposon sequences in the absence of the canonical DNA transposon activity. Despite their relatively small number, they do contribute to the genetic diversity among primate species along with other MEs. Furthermore, the discovery of these retro-DNAs suggests that the L1 TRPT machinery may have been used by more diverse types of RNA transcripts than what we currently know. Future work may include the verification of the retrotransposition activity of these retro-DNAs using *in vitro* and in *vivo* assays and expanding similar analysis to other type of expressive DNA sequences, such as non-coding RNA genes. In addition, investigation into the mechanisms underlying the remaining majority of the diallelic DNA transposons would also be very interesting.

## Acknowledgments

This work is in part supported by grants from the Canadian Research Chair program, Canadian Foundation of Innovation, Ontario Ministry of Research and Innovation, Canadian Natural Science and Engineering Research Council (NSERC), and Brock University to PL, and was made possible by the use of Compute Canada/SHARCNET high-performance computing facilities.

## A list of supplementary tables

Table S1. Composition of diallelic DNA transposons (da-DNAs) by family in the ten primate genomes.

Table S2. The number of retroDNAs with identifiable polyA tails in the 10 primate genomes.

Table S3. Detailed list of retro-DNAs located in genic regions in the 10 primate genomes.

Table S4. Detailed list of retroDNAs and their potential parent sites in the 10 primate genomes.

Table S5. Parent sites associated with genic regions in the ten primate genomes.

Table S6. Expression level of retro-DNAs and potential parent sites in the 7 primate transcriptome.

## A list of supplementary figures

**Figure S1.**
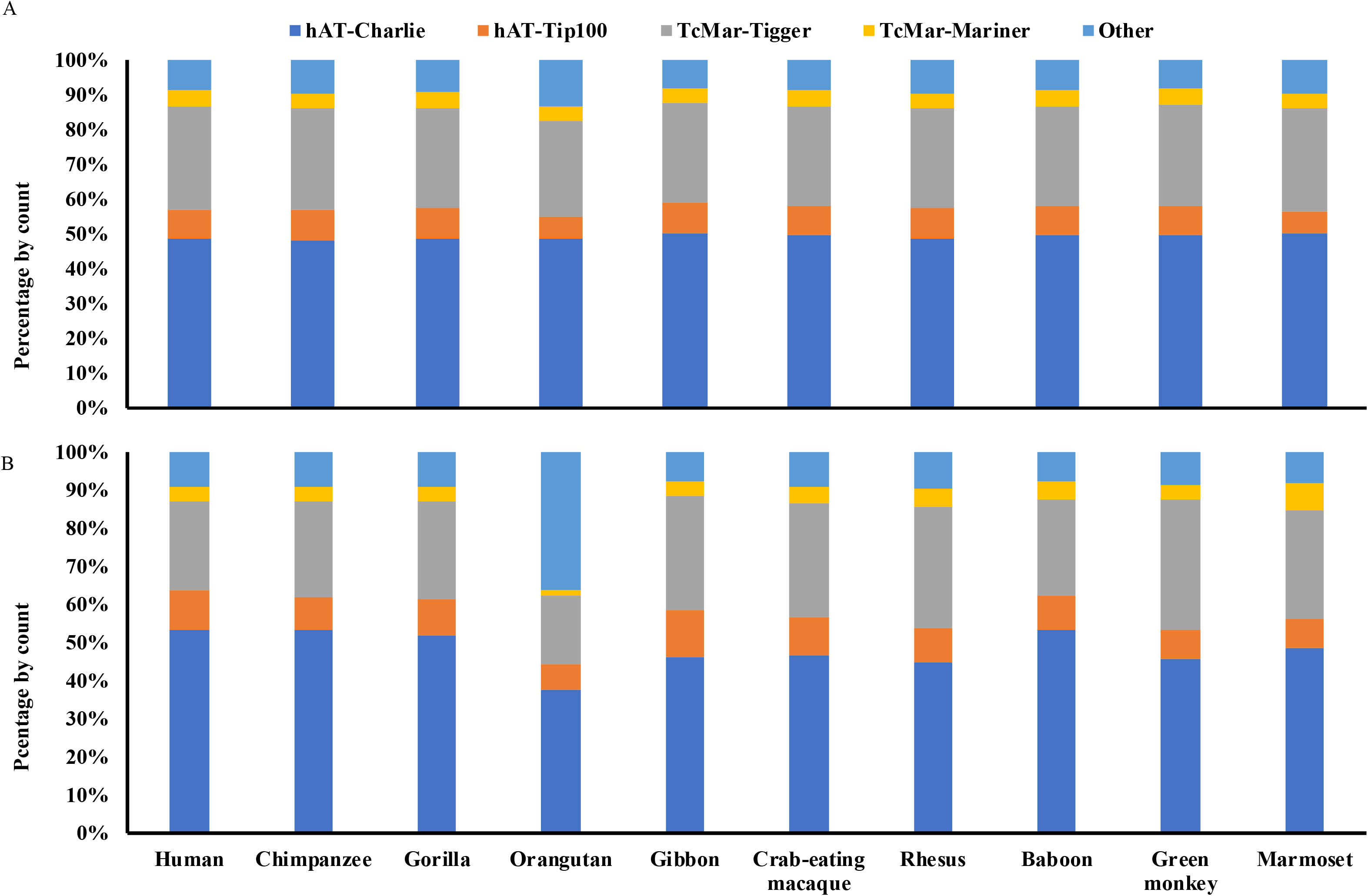
The distribution of diallelic DNA transposons and retro-DNAs by family in the ten primate genomes. Stacked bar plots showing the family of composition of diallelic DNA transposons (A) and retro-DNAs (B) in each of the 10 primate genomes. The color scheme in panel B is the same as in panel A.

**Figure S2.**
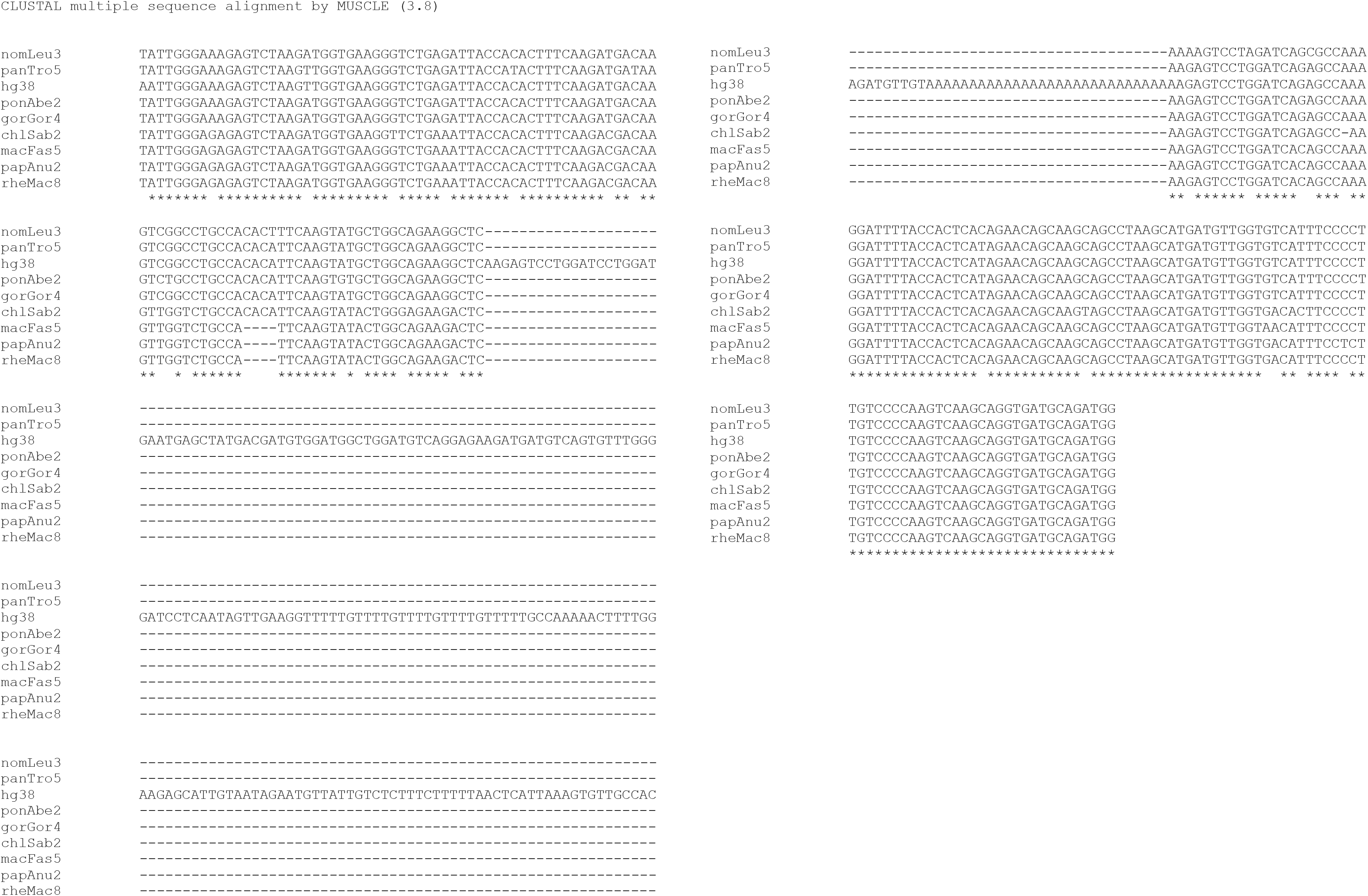
Multiple sequence alignment for the retro-DNA located in the human genome (hg38_chr4:146335052-146335253) and the corresponding pre-integration sequences from eight other primate genomes. The pre-integration sequences from marmoset genome is unavailable likely due to the high level of sequence divergence.

**Figure S3.**
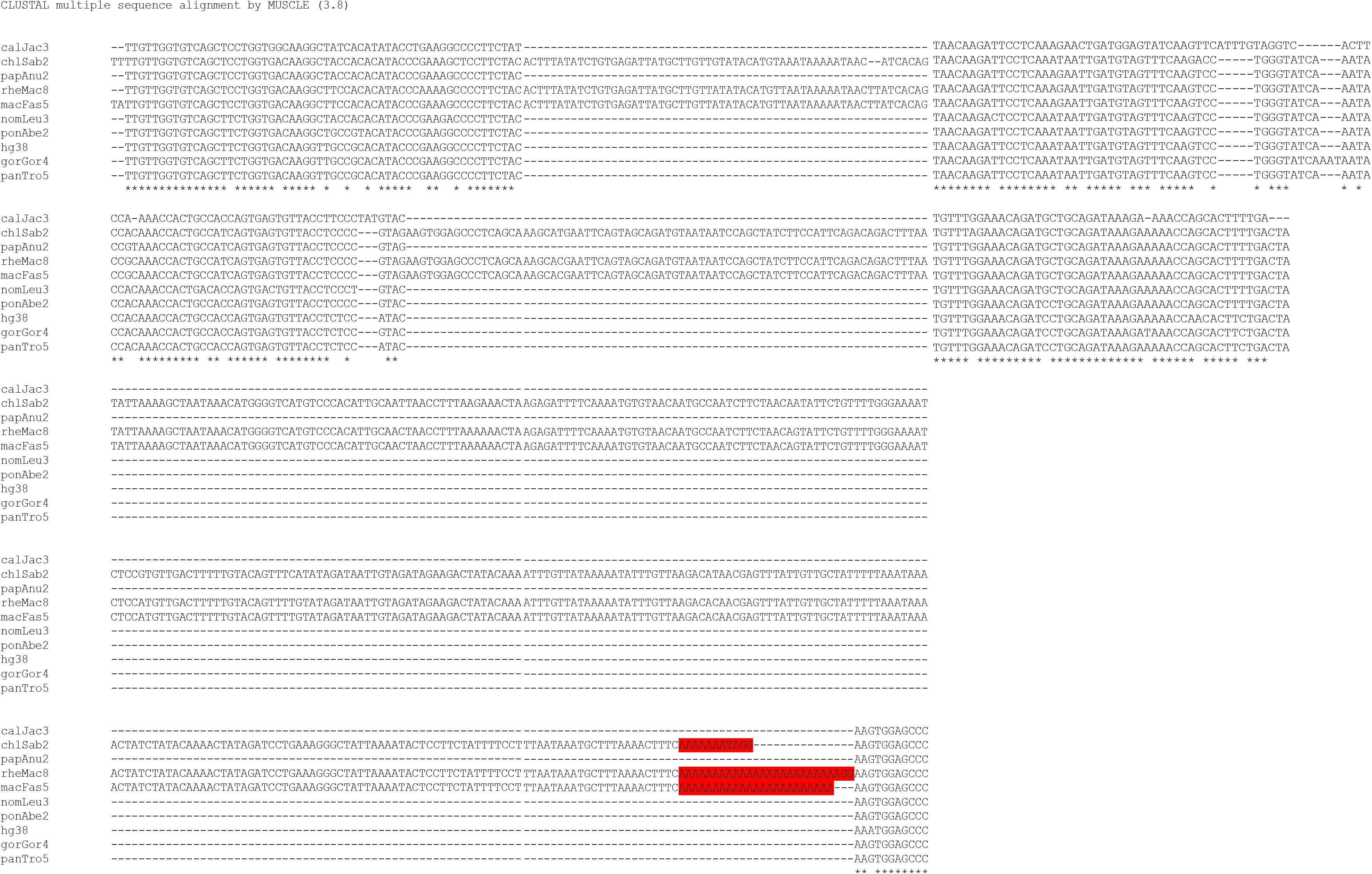
Multiple sequence alignment for retro-DNA located in the green monkey, crab-eating macaque and rhesus genomes (chlSab2_chr8:30005081-30005527/rheMac8_chr8:31992158-31992606/macFas5_chr8:32527581-32528029) and their pre-integration sites from 7 other primate genomes. The red boxes represent possible polyA tail with various length in different genomes.

**Figure S4.**
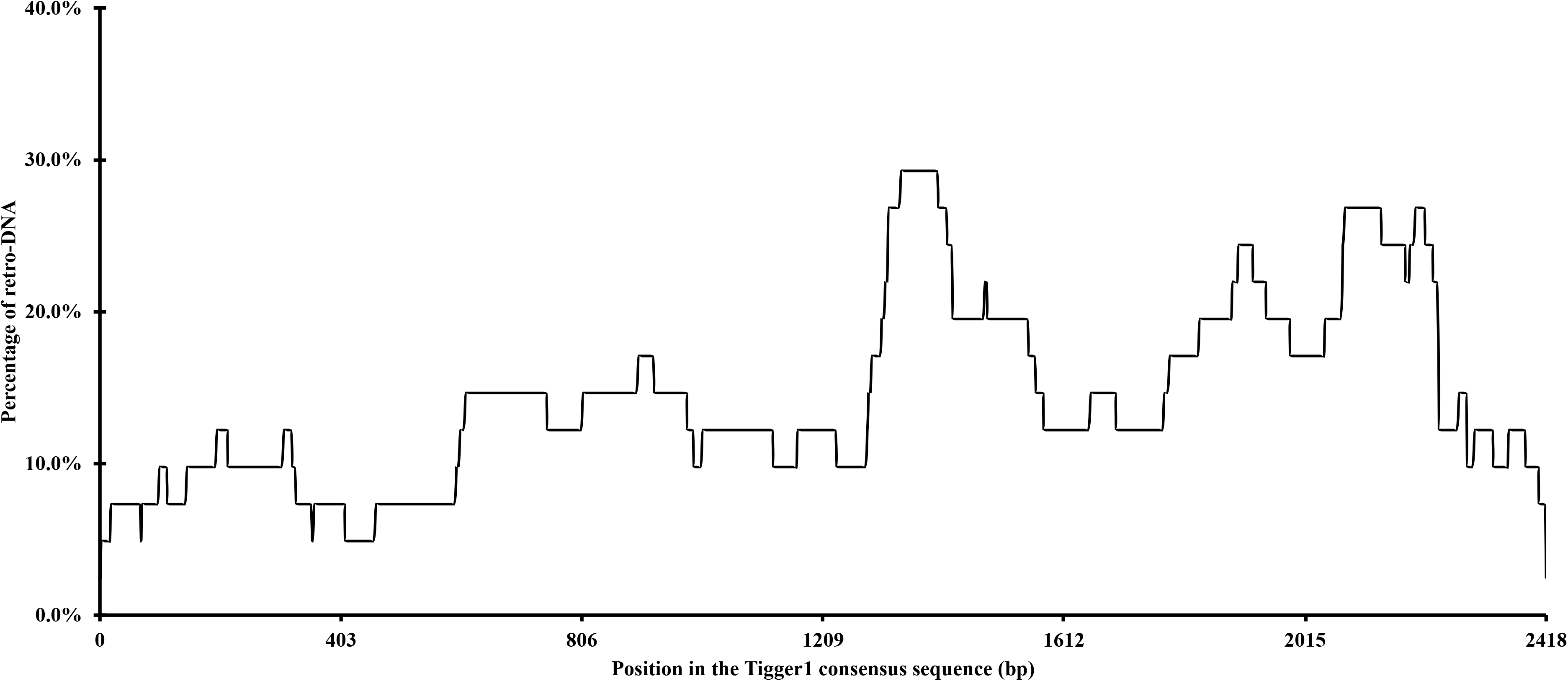
The frequency of the Tigger1 subfamily DNA transposon consensus sequence used for retro-DNA sequences. The plot is based on the data for a total of 41 non-redundant retro-DNA entries from the Tigger1 subfamily.

**Figure S5.**
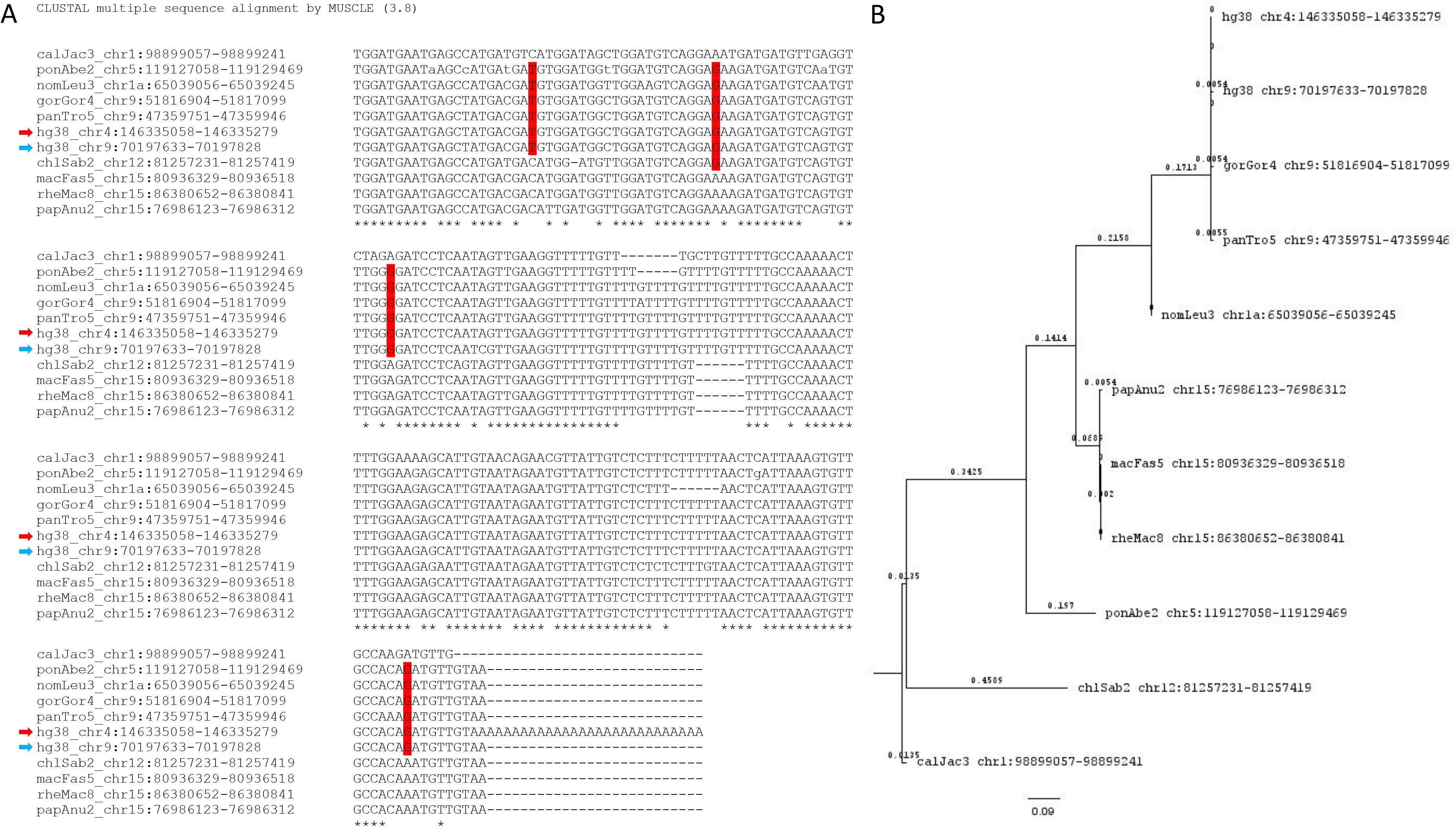
A. Multiple sequence alignment for retro-DNA in the human genome (hg38_chr4:146335052-146335253) and its parent copy (hg38_chr9:70197633-70197828) plus orthologous copies from the other 9 non-human primate genomes. Red arrow indicates the retro-DNA entry, blue arrow indicates the parent copy. SNPs in red boxes are owned by members of the *Hominidae* group. B. Evolutionary analysis by Maximum Likelihood method using the 11 nucleotide sequences from the 10 primate genomes. The evolutionary history was inferred by using the Maximum Likelihood method and Tamura-Nei model. The bootstrap consensus tree inferred from 500 replicates is taken to represent the evolutionary history of the taxa analyzed. Branches corresponding to partitions reproduced in less than 50% bootstrap replicates are collapsed. The percentage of replicate trees in which the associated taxa clustered together in the bootstrap test (500 replicates) are shown next to the branches. Initial tree(s) for the heuristic search were obtained automatically by applying Neighbor-Joining and BioNJ algorithms to a matrix of pairwise distances estimated using the Maximum Composite Likelihood (MCL) approach, and then selecting the topology with superior log likelihood value. This analysis involved 11 nucleotide sequences. There is a total of 222 positions in the final dataset. Evolutionary analyses were conducted in MEGA X.

